# CircPVT1 sponges miR-33a-5p unleashing the c-MYC/GLS1 metabolic axis in breast cancer

**DOI:** 10.1101/2024.06.04.597315

**Authors:** Alina Catalina Palcau, Claudio Pulito, Valentina De Pascale, Luca Casadei, Maria Cristina Valerio, Andrea Sacconi, Daniela Rutigliano, Sara Donzelli, Romana Francesca Auciello, Fulvia Pimpinelli, Paola Muti, Claudio Botti, Sabrina Strano, Giovanni Blandino

**Affiliations:** Microbiology and Virology Unit, San Gallicano Dermatological Institute IRCSS, 00144 Rome, Italy; Translational Oncology Research Unit, IRCCS, Regina Elena National Cancer Institute, Rome, Italy; Department of Chemistry, “Sapienza” University of Rome, Piazzale A. Moro 2, 00185, Rome, Italy; Clinical Trial Center, Biostatistics and Bioinformatics, IRCCS Regina Elena National Cancer Institute, Rome, 00144, Italy; Department of Health Research Methods, Evidence, and Impact, Faculty of Health Sciences, McMaster University, Hamilton, ON, Canada; Department of Biomedical, Surgical and Dental Sciences, University of Milan, Milan, Italy; Department of Surgery, IRCCS Regina Elena National Cancer Institute, Rome 00144, Italy; SAFU Laboratory, IRCCS Regina Elena National Cancer Institute, Rome, Italy

**Author notes:** These two authors contributed equally to this work (Alina Catalina Palcau, Claudio Pulito). Correspondence should be sent to: Giovanni Blandino, M.D., Translational Oncology Research Unit, IRCCS, Regina Elena National Cancer Institute, Rome, Italy., Sabrina Strano, M.D., SAFU Laboratory, IRCCS Regina Elena National Cancer Institute, Rome, Italy.

**Keywords:** non-coding RNAs, metabolism, breast cancer, MYC, Patients Derived Organoids

## Abstract

Altered metabolism is one of the cancer hallmarks. The role of circRNAs in cancer metabolism is still unexplored. Herein, we initially found that the expression of circPVT1 was significantly higher in tumoral tissues than in non-tumoral breast tissues. Basal like breast cancer patients with higher levels of circPVT1 exhibited shorter disease-free survival compared to those with lower expression. CircPVT1 ectopic expression rendered fully transformed MCF-10A immortalized breast cells and increased tumorigenicity of TNBC cell lines. Depletion of endogenous circPVT1 reduced tumorigenicity of SUM-159PT and MDA-MB-468 cells. 1H-NMR spectroscopy metabolic profiling of circPVT1 depleted breast cancer cell lines revealed reduced glycolysis and glutaminolitic fluxes. Conversely, MCF-10A cells stably overexpressing circPVT1 exhibited increased glutaminolysis. Mechanistically, circPVT1 sponges miR-33a-5p, a well know metabolic microRNA, which in turn releases c-MYC activity which promotes transcriptionally glutaminase, which converts glutamine to glutamate. CircPVT1 depletion synergizes with GLS1 inhibitors BPTES or CB839 to reduce cell viability of breast cancer cell lines and breast cancer-derived organoids. In aggregate, our findings unveil the circPVT1/miR-33a-5p/Myc/GLS1 axis as a pro-tumorigenic metabolic event sustaining breast cancer transformation with potential therapeutic implications.

## Introduction

Breast cancer is one of the most frequent causes of death among women worldwide. GLOBOCAN estimated that female breast cancer overcame lung cancer as the leading cause of global cancer incidence in 2020 with 2.3 million new cases [1]. Breast cancer is a rather heterogeneous type of tumor that can be molecularly classified in four different subtypes: Luminal A, Luminal B, HER2-positive and Triple Negative breast cancer. Among the different subtypes, triple negative breast cancer that express no oestrogen, progesterone and human epidermal growth factor receptor 2 is the most aggressive one with the poorest prognosis. Indeed, in the absence of a specific target therapy, the common treatment is based on chemotherapy or radiotherapy [2].

CircRNAs have been proposed initially as splicing errors but actually they play important role within cells. These RNAs molecules due to their covalently closed structure are very resistant in body fluids where they can be evaluated through liquid biopsy [3]. Indeed, their expression it has been demonstrated to be altered in tumors such as breast cancer, in which they can either prevent or trigger tumorigenesis [1, 4, 5]. Circular RNAs function as scaffolds, as they interact directly with proteins or also be translated in a cap-independent mechanism generating small functional peptides [6]. The most studied and also debated mechanism of circRNA function is the miRNAs sponge activity. miRNAs are small sequences of 22-24 nucleotides playing an important role in the regulation of gene expression by binding to the 3’UTR of their mRNAs targets [7]. Circulating miRNAs seem to be promising biomarkers for diagnosis and prognosis of diseases including cancer [8, 9]. miRNAs have been proposed as potential biomarkers since their expression levels can be measured also in body fluids in order to evaluate the progress of the therapy or even to instruct toward a specific therapy [10]. CircPVT1 originates from the circularization of exon 2 of PVT1 gene, located near MYC locus, that encodes also for a lncRNA called lncPVT1 [11, 12]. It has been shown that circPVT1 exerts oncogenic activities in several tumors including gastric cancer or head and neck squamous cell carcinoma [13, 14]. Altered metabolism is one of the pivotal hallmarks of cancer [15]. Cancer cells require high amount of energy in terms of ATP to sustain their growth but also to obtain precursors of important molecules. Several metabolic pathways become altered during transformation including glycolysis or glutaminolysis. Glutamine serves as a source of reduced nitrogen for biosynthetic reactions and also as source of carbon for molecules biosynthesis [16]. Glutaminase, the key enzyme of glutaminolysis converts glutamine into glutamate which in turn is converted in α-ketoglutarate (α-KG) by glutamate dehydrogenase-1 (GLUD1), allowing the entrance in the tricarboxylic acid (TCA) cycle. Recently, altered glutamine metabolism has been associated with TNBC progression and metastasis [17].

In the present study, we found that aberrantly expressed circPVT1 sponges miR-33a-5p and unleashes MYC/GLS metabolic axis sustaining altered glutaminolysis. Interestingly, the treatment of triple negative breast cancer-derived organoids with GLS1 inhibitors such as CB839 and BPTES strongly reduced cell viability. Altogether these findings with those documenting circPVT1 aberrant expression in breast cancer patients associated with poor overall survival, identify an unexplored network which might hold therapeutic potential.

## Methods

### Cell culture and transfection

Human breast cancer cell lines SUM-159PT, MDA-MB-468 and normal human breast epithelial cell line MCF-10A were purchased from the American Type Culture Collection (ATCC, Manassas, VA, USA). SUM-159PT and MDA-MB-468 cells were grown in DMEM/F12 Glutamax medium (Invitrogen, Carlsbad, CA, USA) supplemented with 10% fetal bovine serum, 100 units/mL Pen/Strep antibiotic and Insulin 5 μg ml−1 (Sigma, Saint Louis, USA) at 37 °C in a balanced air humidified incubator with 5% CO2. MCF-10A cells were grown in DMEM/F12 Glutamax (Invitrogen, Carlsbad, CA, USA) supplemented with 10% horse serum and 100 μL of EGF 20 ng/ml, 500 μL of Antibiotic 100X, 500 μL of HC 500 ng/ml and 500 μL of Human insulin 0.01 mg/ml at 37 °C in a balanced air humidified incubator with 5% CO2.

Lipofectamine RNAimax (Invitrogen, Carlsbad, CA, USA) was used in accordance with the manufacturer’s instruction for transfection with siRNAs and miRNA mimics. SiRNAs were used at the final amount of 300 pmol in 100 mm dish. si-circPVT1 5’-CUUGAGGCCUGAUCUUUUA-3’ was used for functional in vitro experiments. For mature miR-33a-5p overexpression, we used the mirVana miR-33a-5p mimic (Ambion) at a final concentration of 5 nM and as control we used the mirVana miRNA mimic, Negative Control #1 (Ambion), at the same concentration. The circPVT1 overexpression in MCF-10A cells was performed using 4 μg pcDNA3-circPVT1[14] and 4 μg pcDNA3 vector as control. Plasmids were transfected with Lipofectamine 2000 (Invitrogen Carlsbad, CA, USA) in accordance with the manufacturer’s instruction at a final concentration of 1 μg in a 60 mm dish. Cells were collected 48–72 h post transfection for subsequent analyses.

### RNA processing and qPCR

The total RNA was extracted with TRizol (Thermo Fisher Scientific, Rockford, IL, USA) following the manufacturer’s instructions and the concentration, purity, and quality of total RNA were assessed using a Nanodrop TM 1000 spectrophotometer (Nanodrop Technologies).

cDNA synthesis and qRT-PCR

One microgram of total RNA was reverse transcribed at 37□°C for 60□min in the presence of random hexamers and Moloney murine leukaemia virus reverse transcriptase (Invitrogen, Carlsbad, CA, USA). Specific oligonucleotide primers for

ACTIN Fw: 5′-GGCATGGGTCAGAAGGATT-3′ and

Rv: 5′-CACACGCAGCTCATTGTAGAAG-3′.

CircPVT1 Fw:5’CGACTCTTCCTGGTGAAGCATCTGAT-3’ and

Rv:3’ TACTTGAACGAAGCTCCATGCAGC-5’;

C-MYC Fw: 5′-CTCCTGGCAAAAGGTCAGAG-3’ and

Rv: 5′-TCGGTTGTTGCTGATCTGTC-3′;

GLS Fw: 5’-TTCCAGAAGGCACAGACATGGTTG-3’ and

Rv: 5’-GCCAGTGTCGCAGCCATCAC-3’ were used for PCR analyses. Gene expression levels were measured by quantitative real-time PCR using the SYBR Green assay (Thermo Scientific, Rockford, IL, USA)on a StepOne instrument (Thermo Scientific, Rockford, IL, USA). Small amounts of RNA (10□ng) were reverse transcribed using the TaqMan microRNA Reverse Transcription Kit (Applied Biosystems) in a final volume of 10□µl using an ABI Prism 7000 Sequence Detection System (Thermo Scientific, Rockford, IL, USA). The PCR reactions were initiated with a 10-min incubation at 95□°C followed by 40 cycles of 95□°C for 15□s and 60□°C for 60□s. qPCR quantification of miRNA expression was performed using TaqMan MicroRNA® Assays (Thermo Scientific, Rockford, IL, USA) according to the manufacturer’s protocol. RNU48 and RNU44 were used as an endogenous control to normalize miRNA expression. All reactions were performed in triplicate. For circPVT1, PVT1, MYC, GAPDH and GLS1 gene expression analysis, reverse transcription and RT-qPCR were performed using MMLV RT (Invitrogen, Carlsbad, CA, USA) and SYBR Green® Assays (Thermo Scientific, Rockford, IL, USA), respectively, according to the manufacturers’ instructions.

### PDO cultures

For organoid generation, tissues were processed as described in Sash et al., 2018 [18]. Briefly, tissues were mechanically processed until small portions (1mm3) were obtained. Then they were enzymatically digested for 1 h at 37°C with Tumor Dissociation Kit, human (Miltenyi, Bergisch Glabach, Germany) according to the manufacturer’s instructions. The suspension was strained over a 70 μm filter. Isolated cell clusters were resuspended in Matrigel® (Corning, New York, USA) and plated in 24-well plates.

PDO were grown in Ad-DM medium supplemented described as in Donzelli S. and colleagues [19] with 1X Glutamax (Thermo Scientific, Rockford, IL, USA), 10mM Hepes (Thermo Scientific, Rockford, IL, USA), 1X Penicillin/Streptomycin (Thermo Scientific, Rockford, IL, USA), 50 μg/mL Primocin (InVivogen Toulouse, France), 1X B27 supplement (Thermo Scientific, Rockford, IL, USA), 250 ng/mL R-spondin 1 (PeproTech, Cranbury, NJ, USA), 5 nM Heregulin β−1 (PeproTech, Cranbury, NJ, USA), 100 ng/mL Noggin (PeproTech, Cranbury, NJ, USA), 20 ng/mL FGF-10 (PeproTech, Cranbury, NJ, USA), 5 ng/mL FGF-7(PeproTech, Cranbury, NJ, USA), 5 ng/mL EGF (PeproTech, Cranbury, NJ, USA), 5 μM A83-01 (Tocris Bioscience), and 500 nM SB202190 (PeproTech, Cranbury, NJ, USA). Moreover, 5 μM Y-27632 (PeproTech, Cranbury, NJ, USA) was added to culture media for the first three days of culture. When confluency was reached, organoids were dissociated by resuspension in 2 mL TrypLE Express (Invitrogen Carlsbad, CA, USA) and incubation for 10 min at room temperature. After enzyme neutralization and washing, organoids were resuspended in Matrigel and reseeded as above in order to allow the formation of new organoids.

Bright-field imaging of organoids was performed on an NEXCOPE microscope.

Generation of patient-derived organoids from breast cancer was approved by institutional review board of Regina Elena National Cancer Institute and appropriate regulatory authorities (approval no. IFO 1270/19). All patients signed an informed consent.

### Cell and PDOs viability assay

Viability of treated cells and PDOs was assessed using ATPlite assay (Perkin Elmer, Massachusset, USA) accordingly to the manufacturer’s instructions. Cells (8×102cells) and PDOs were seeded in 96 well-plates and cultured for 24hrs and treated for 72 hrs with BPTES or CB839. Each plate was evaluated immediately on a microplate reader (EnSpire Technology, Perkin Elmer, Massachusetts, USA).

### Opera Phoenix plus

The PDOs were dissociated in 2ml of TrypleE (Invitrogen 12605036), incubated at 37° for 10 minutes. After digestion, 10mL of medium is added and cells were centrifugated for 5 minutes at 1000 rpm. The dissociated PDOs were, then, resuspended in 700uL of Matrigel and 1000 cells per well were seeded (Matribot, Corning) in a PhenoPlate 96-well, black, optically clear flat-bottom, 96 wells (Revvity, Waltham, Massachusets,USA). After drops solidification, 100 μl of medium is added. When PDOs reached an area dimension of more than 500 μm2 they were treated. Number, area, perimeter, width, length and cytotox staining (Incucyte Cytotox Green Dye, Sartorius) of the spheroids were assessed by using the Opera Phenix® Plus high throughput microplate confocal imager (Revvity) and calculated by using the Harmony High-Content Imaging and Analysis Software (Revvity).

### Clonogenic assay

Transfected BC cells at a density of 1,000 were seeded in six-well plates. Cell colonies were subsequently washed, fixed, and stained until the colonies were visible. Then, colonies were counted and imaged.

### Transwell invasion assay

Transfected BC cells were added into the upper chamber with 200 μL of serum-free medium. After culturing for 24 h for the SUM-159PT, MDA-MB-468 and MCF-10A cell lines, cells that migrated to the opposite side of the filter were fixed, stained, imaged (Leica Microsystems, Germany), and counted.

### Protein extracts and western blot analysis

Cells were homogenized on ice for 30 min in a lysis buffer composed by 50 mM, Hepes pH 7.5, 5 mM EDTA pH 8.0, 10 mM MgCl2, 150 mM NaCl, 50 mM NaF, 20 mM β-glicerophosphate, 0.5% NP40, 0.1 mM sodium orthovanadate, 1 mM PMSF, 1 mM dithiothreitol (DTT), and protease inhibitor cocktail (Roche). Lysates were clarified by centrifugation for 10 min, max speed, at 4°C. Proteins (30 μg/lane) were separated on 10% SDS-polyacrylamide gels and transferred to nitrocellulose membranes. Immunoblots were probed with the following primary antibodies: rabbit monoclonal anti-c-Myc (DO1; Oncogene Science Uniondale, NY, USA), and mouse monoclonal anti-GAPDH (Calbiochem). Immunostained bands were detected by a chemiluminescent UVITEC Alliance 4.7 instrument (Cambridge, UK). ECL solution (entry-level peroxidase substrate for enhance chemiluminescence) (Thermo Scientific, Rockford, IL, USA) was loaded on the membrane in order to allow the chemiluminescent reaction between horseradish peroxidase (HRP) labelled on the secondary antibody and the peroxidase substrate of ECL solution. The reaction generates energy that is released in the form of light and in this way the protein signal is detected by the camera.

### Chromatin Immunoprecipitation (ChIp)

ChIP Assay Kit (Millipore, Bedford, MA) was used according to manufacturer’s instructions. In brief, the 1% formaldehyde cross-linked chromatin was sonicated into fragments and then immunoprecipitated using MYC and POL2(ser5p) antibodies. IgG was used as negative control. DNA fraction was analyzed by qRT-PCR.

### MagIC Beads RNA pull down

Samples of total, unfragmented RNA from SUM-159PT and MCF-10A #7 cells were incubated with MagIC Beads targeting human circPVT1 transcript according to manufacturer’s instructions (https://elementzero.bio/magic-beads-rna-enrichment/). RNA attached to the beads was washed, eluted and subjected to cDNA synthesis. Levels of miR-33a-5p, miR-145 and miR-203 were measured in the input and enriched samples with RT-qPCR.

### Subcellular fractionation

Nuclear and cytoplasmic extraction reagents (Thermo Fisher Scientific, Rockford, IL, USA) were used for subcellular fractionation of BC cells. We used H3 as the nuclear control and Tubulin as the cytoplasmic control.

### Analysis of GLS promoter

Lasagna 2.0 web-tool to analyse GLS promoter. The promoter sequences are related to the human genome GRch38/hg38.

### Seahorse analysis

Analyses of Glutamine Dependency were performed on live cells using a Seahorse XF HS Mini Bioanalyser (Agilent Technologies; Santa Clara, CA, USA) and the Mito Fuel Flex Test (Agilent Technologies; Santa Clara, CA, USA). SUM-159PT and MCF-10A #7 cells overexpressing miR-33a-5p (4 × 10^4^ cells/well) were seeded and grown over night prior to analysis. Assays were performed using protocols suggested by the manufacturer (Agilent Technologies; Santa Clara, CA, USA). Briefly, after a 30-min calibration of the XF sensor with a preincubated sensor cartridge, the cell plate was loaded into the analyzer, and Oxygen Consumption Rate (OCR) was analyzed under basal conditions. Glutamine Dependency was tested by first injecting BPTES (3 μM), an inhibitor of allosteric GLS1 glutaminase, followed by inhibition of carnitine palmitoyl-transferase 1A (CPT1A) and mitochondrial pyruvate transporter (MPC) using Etomoxir (4 μM) and UK5099 (2 μM) respectively. OCR was measured after BPTES injection and following the injection of the other two inhibitors (All Inhibitors). After performing MFFT assays, cells were stained with DAPI and the cell count of each well was determined by imaging the cells using Cell Profiler. Once the assay was normalized on cell number, Glutamine Dependency was calculated with the following equation:

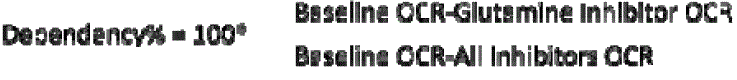

### 1H-NMR spectroscopy

All 2D 1H J-resolved (JRES) NMR spectra were acquired on a 500LMHz VNMRS Varian/Agilent spectrometer (Agilent, Santa Clara, CA) at 25 °C using a double spin echo sequence with pre-saturation for water suppression and 16 transients per increment for a total of 32 increments. These were collected into 16Lk data points using spectral widths of 8 kHz in F2 and 64LHz in F1. Each free induction decay (FID) was Fourier transformed after a multiplication with sine-bell window functions in both dimensions. JRES spectra were tilted by 45°, symmetrized about F1, referenced to lactic acid at δHL=L1.33Lppm and the proton-decoupled skyline projections (p-JRES) exported using Agilent VNMRJ 3.2 software. The exported p-JRES were aligned, corrected for baseline offset and then reduced into spectral bins with widths ranging from 0.02 to 0.06□ppm by using the ACD intelligent bucketing method (1D NMR Manager software, ACD/Labs, Toronto, Canada). This method sets the bucket divisions at local minima (within the spectra) to ensure that each resonance is in the same bin throughout all spectra. The area within each spectral bin was integrated and in order to compare the spectra, the integrals derived from the bucketing procedure were normalized to the total integral region. Metabolites were identified using an in-house NMR database and literature data and confirmed by 2D homo– and hetero-nuclear NMR spectroscopy.

### NMR spectra pre-processing treatment

The 1D skyline projections exported were aligned and then reduced into spectral bins with ranging from 0.01 to 0.02□ppm by using the ACD intelligent bucketing method (1D NMR Manager software (ACD/Labs, Toronto, Canada). To compare the spectra, the integrals derived from the binning procedure were normalized to the total integral region, following exclusion of bins representing the residual water peak (4.33–5.17□ppm) and the TSP peak (0.5–0.5□ppm). The resulting data was used as input for multivariate analysis: Principal Component Analysis (PCA and Orthogonal projections to latent structures discriminant analysis (OPLS-DA) were performed using SIMCA-P□+□version 12 (Umetrics, Umea, Sweden).

### Statistical analysis

The resulting data was used as input for univariate and multivariate analysis PCA34 and OPLS-DA35. PCA and OPLS-DA were conducted using SIMCA-P+ version 12 (Umetrics, Umea, Sweden). For microRNA analysis, Pearson’s correlation coefficient was calculated to assess quality of replicates. Generally, Student’s t-test was used to assess significance of the data and P-values ≤0.05 were considered statistically significant.

## Results

### Aberrant expression of circPVT1 elicits pro-tumorigenic effects in breast cancer cells

We found that circPVT1 expression was significantly higher in breast cancer patient tissues when compared with non-tumoral ones in The Cancer Genome Atlas (TCGA) data set (Fig. 1a). CircPVT1 aberrant expression was more pronounced in advanced stages (III/IV) compared to I/II stages (Fig. 1b). Interestingly, basal like breast cancer patients expressing higher levels of circPVT1 exhibit shorter overall free survival (Fig. 1c). dPCR analysis using specific primers spanning the junction of circPVT1 circularization revealed that circPVT1 expression was higher in breast cancer tissues than in matched non-tumoral tissues derived from patient enrolled consecutively at the IRCCS Regina Elena National Cancer Institute (Fig.1d). To ascertain further circPVT1 oncogenic role in breast cancer, we found that ectopically expression of circPVT1 promoted colony formation ability and migration of two SUM-159PT and MDA-MD-468 breast cancer cell lines (Fig. 1e-f) (Fig. S1a-b). Congruently, depletion of circPVT1 caused a reduction of colony formation and migration of both SUM-159PT and MDA-MD-468 cells (Fig. 1g-h) (Fig. S1a-b). Interestingly, 1H-NMR spectroscopy metabolic profiling of culture media of circPVT1 depleted SUM-159PT cells (SUM_sicircPVT1) compared to that of control cells (SUM_siSCR) was performed. Firstly, 1H-NMR data were analysed by using unsupervised PCA analysis (data not shown). Secondly, the same 1H-NMR dataset was analysed with the supervised method of OPLS-DA. A good predictive model (Q2=0.89) with one predictive and sic orthogonal LVs R2X=76% and R2Y=100% was obtained (Fig. 1i). A t-test applied on the predicted LV1 confirmed significant metabolic difference between SUM_sicircPVT1 and SUM_siSCR cells (p < 0.00001). The metabolic differences are expressed as fold change over the control-silenced cells (Fig. 1j). By looking at the main metabolites correlated to a tumor phenotype, we observed that SUM_sicircPVT1 cells exhibited higher concentration of glucose and glutamine as well as lower concentrations of glutamate, lactate, alanine and acetate. These findings might indicate decreased glycolytic and glutaminolitic fluxes and increased oxidative metabolism, thereby suggesting that cancer cells produce less energy needed to grow and spread (Suppl. Fig.1c). To further understand whether the depletion of circPVT1 reprograms metabolism toward a less tumorigenic phenotype we compared its effect with that induced by metformin, an oral anti-diabetic drug with well-known metabolic activities [20, 21]. Interestingly, we found that metformin treatment downregulated circPVT1 expression in breast cancer cells (Fig. 1k). We then performed 1H-NMR spectroscopy metabolic profiling of culture media derived from SUM_siSCR treated with metformin and SUM_sicircPVT1 cells. SUM_siSCR untreated cells were included as control. The derived NMR profiles were compared by using OPLS-DA obtaining a good predictive model (Q2=0.9) with one predictive and three orthogonal LVS with R2X=86% and R2Y=100% (Fig. 1l). A t-test applied on LV1 showed statistically significant metabolic differences between SUM_siSCR treated with metformin and SUM_sicircPVT1 cells thereby highlighting a broader metabolic impact of metformin than circPVT1 depletion on breast cancer cells (Fig. 1m). Interestingly on the LV2 we evidenced only statistically significant metabolic differences when SUM_siSCR untreated control cells were compared to both SUM_siSCR treated with metformin and SUM_sicircPVT1 cells (Fig. 1m). The loading analysis revealed that LV1 is indicative of glycolysis pathway while LV2 of the fatty acid synthesis. The quantitative OPLS-DA approach showed that the impact of circPVT1 depletion on fatty acid synthesis was greater than metformin treatment (LV2), while that on the glycolysis pathway (LV1) was more evident on metformin-treated breast cells (Fig. 1l-m). In aggregate, our findings report that aberrant expression of circPVT1 exerts pro-proliferative and pro-migratory effects sustaining altered glutaminolysis.

### Ectopic expression of circPVT1 promotes aberrantly glutaminolysis in the non-tumorigenic cell line MCF-10A breast cell line

To further ascertain the pro-tumorigenic activity of circPVT1 in breast cancer, we stably expressed circPVT1 into a non-tumorigenic breast cell line, MCF-10A that exhibits endogenous circPVT1 levels lower than TNBC cell lines (Fig. 2a) (Fig. S1d). Intriguingly, MCF-10A stable expressing high levels of circPVT1 were, indeed, able to form colonies and to migrate compared to control cells, thereby strengthening the oncogenic role of circPVT1 in breast cancer (Fig. 2b-c). Subsequently, 1H-NMR spectroscopy metabolic profiling of culture media of three stably overexpressing circPVT1 MCF-10A (#2, #4, #7) clones compared to control cells was performed. Principal Component Analysis and OPLS-DA were applied to MCF-10A-circPVT1#7 that positioned the furthest among circPVT1 clones when compared to control cells (Fig. 2d-e). Congruently we built an OPLS-DA model containing only clone #7 (Fig. 2f). A t-test applied on the predictive LV1 confirmed significant metabolic differences between the two groups (p < 0.00001). Loading analysis revealed that the flux of the glutaminolytic pathway was increased in MCF-10A-circPVT1#7 compared to control cells (Fig. 2g). Altogether these findings data unveil an oncogenic role of circPVT1 in breast cancer which leads to altered glutaminolysis.

### CircPVT1 sponges the metabolic miR-33a-5p in breast cancer cells

To mechanistically decipher the contribution of altered circPVT1 expression to cancer metabolism of breast cancer cells we aimed to assess its ability to sponge microRNAs with well-established metabolic activities. We have previously reported that metformin-induced metabolic reprogramming of breast cancer cell lines was at least partially, mediated by the up-regulation of miR-33a-5p [22] and the downregulation of miR-21-5p [23]. Interestingly we found that circPVT1 may interact by direct binding with miR-33a-5p (Fig. 3a upper panel) and also that MCF-10A-circPVT1 expressing clones exhibited reduced expression of miR-33a-5p compared to control cells (Fig. 3a lower panel). Subcellular localization of circPVT1 revealed its predominant localization in the cytoplasmic fraction (Fig. 3b). Depletion of circPVT1 in SUM-159PT cells led to increased miR-33a-5p expression (Fig. 3c). Unlike miR-33a-5p, no modulation of miR-21-5p was evidenced (Fig. 3c). Altogether these findings prompted us to assess whether circPVT1 sponges miR-33a-5p in breast cancer cells lines. RNA immunoprecipitation analysis by using beads produced through a direct synthesis of circPVT1-capture probes on magnetic nanoparticles (see Method section) was performed in both SUM-159PT and in MCF-10A clone #7 breast cancer cells. Notably, a direct biding of miR-33a-5p to circPVT1 was evidenced in both cell lines (Fig. 3d). This binding was not evidenced for miR-21-5p which was used as unrelated miRNA to test binding specific of circPVT1 to miR-33a-5p (Fig. S1i). Functionally, we found that ectopic expression of miR-33a-5p rendered SUM-159PT and MCF-10A/circPVT1#7 cells more dependent from glutamine metabolism than control cells thereby rescuing at least partially the aberrant of circPVT1 on glutaminolysis (Fig. 3e-f). Collectively our findings originally show that circPVT1 sponging the metabolic miR-33a-5p contributes to altered glutaminolysis.

### c-MYC mediates circPVT1/miR-33a-5p-induced metabolic alteration in breast cancer cells

To further dissect the role of circPVT1 in altered glutaminolysis of breast cancer cells we focused on c-MYC, a well-known miR-33a-5p target with reported transcriptional activity on the regulation of key metabolic enzymes. Indeed, it has been reported that c-MYC fine tunes glutaminase (GLS) translation, the enzyme converting glutamine to glutamate, by directly suppressing miR-23a and miR-23b expression, thereby leading to aberrant GLS expression [24]. We have previously shown data that miR-33a-5p directly binds the c-Myc 3’-UTR abrogating c-MYC expression in breast cancer cells [22]. Interestingly we found a direct correlation when comparing circPVT1 and c-MYC expression levels in breast cancer TGCA database and MCF-10A#circPVT1 expressing clones (Fig. 4a-b). Congruently, c-MYC protein expression was increased in MCF-10A#circPVT1 expressing clones and reduced in SUM-159PT cells upon depletion of circPVT1 (Fig. 4c-d). In order to understand the role of c-Myc in the circPVT1-mediated induction of glutaminolysis, we analysed the metabolites released in the media of MCF-10A circPVT1 clone #7 silenced for c-Myc expression (Fig. 4e-f). The medium samples derived from the control cells (M10_siSCR clone#7) and those deriving from the c-Myc-depleted cells (Si-Myc clone#7) were analysed by using 1H-NMR spectroscopy. OPLS-DA analysis was used to compare NMR profiles (Fig. 4g). A t-test applied on LV1 showed significant metabolic differences between the medium samples of the two groups (p < 0.00001) (Fig. 4g). Interestingly, c-MYC depleted cells exhibited increased glucose and glutamine consumption thereby indicating when compared to control cells. These findings suggest increased oxidative metabolism following c-MYC depletion in breast cancer cells overexpressing circPVT1 (Fig. 4h).

### c-MYC regulates transcriptionally GLS1 expression

We aimed to assess whether c-MYC could transcriptionally regulate the expression of GLS. A positive correlation between c-MYC and GLS expression was evidenced in both breast cancer TGCA database and in MCF-10A cell clones overexpressing circPVT1 (Fig. 5a-b). Interestingly c-MYC depletion in cell overexpressing circPVT1 significantly reduced GLS1 transcript (Fig. 5c-d). By using Lasagna software, we identified a putative binding site for c-Myc on GLS promoter (4850-4811 upstream GLS transcriptional starting site) (Fig. 5e). Furthermore, the analysis of c-Myc ChIP-seq data deposited on the CistromeDataBase (http://cistrome.org/) revealed a binding of the c-Myc protein onto a specific binding site of GLS promoter region in MCF-10A cells (Fig. 5f). Congruently, chromatin immunoprecipitation assays confirmed a direct binding of c-Myc on GLS promoter region both in SUM-159PT and in MCF-10A clones overexpressing circPVT1 (Fig. 5g-h). The same promoter region was enriched in active Polymerase II (ser5p) thereby indicating the active transcription of GLS gene associated to the c-Myc protein binding (Fig. 5g-h). Altogether these findings support the critical role of c-MYC transcriptional activity in the aberrant regulation of GLS as downstream effector of an oncogenic cascade instigated by the abnormal expression of circPVT1 in breast cancer cells.

### CircPVT1 depletion sensitizes breast cancer cells and TNBC derived organoids to the glutaminase inhibitors BPTES and CB839

Growing evidence has shown that glutamine addiction in breast cancer cells might represent a novel therapeutic target. Indeed, inhibition of glutaminase through specific inhibitors is sought to prevent aberrant cell proliferation and sensitize cancer cells to anticancer treatment. We found that either metformin-induced downregulation (Fig. S2a) or siRNA-mediated (Fig. 6a and Fig. S2b) depletion of circPVT1 expression rendered SUM-159PT and MDA-MB-468 more sensitive to the killing induced by BPTES and CB839 glutaminase inhibitors (Fig. 6b and Fig. S2c). Interestingly, MCF-10A cells stably overexpressing circPVT1 are less prone to the BPTES and CB839-induced cell killing effects than control cells (Fig. 6c and Fig. S2d). Breast tumor organoids derived from three independent triple negative breast cancer patients (T3N0 grade 3, T2N3 grade 3, T4N2a grade 3) were assessed for endogenous expression of circPVT1, miR-33a-5p, MYC and GLS which resulted comparable to that of the originating breast cancer tissues (Fig. 6d). Interestingly ectopic expression of miR-33a-5p led to downregulation of c-Myc and consequently of GLS1 expression levels, thereby providing strong evidence to the suitability of TNBC-derived PDOs to assess the response to glutaminase inhibitors (Fig. 6e). Indeed, BPTES treatment provoked a dose-dependent effect on three independent breast cancer patient derived organoids cell viability highlighting the critical role of glutaminase in the aberrant survival of breast cancer cells (Fig. 6f). To further assess the impact of glutaminase inhibitor on TNBC-derived PDOs we performed high-content imaging analysis on one representative PDO #240 with the aid of Opera Phenix Plus platform. This allowed evaluating the effect of BPTES at single cell organoid resolution. As shown in (Fig. 6g-h) the number of cytotox positive organoid structures as sign of death increased upon BPTES treatment. Similarly, the ratio of death vs live organoids and ATP metabolic activity increased upon BPTES treatment (Fig. 6i-j). CB839 treatment induced similar effects but to a lesser extent than BPTES (Fig. 6k).

Collectively, these findings indicate that aberrant expression of circPVT1 might contribute to glutamine addiction of breast cancer cells leading to the aberrant activation of glutaminase. Herein we also provide insights supporting the role of selective glutaminase inhibitors as anticancer therapy intercepting oncogenic networks initiated by aberrant expression of circular RNAs such as circPVT1 in breast cancer.

## Discussion

Altered metabolism is a cancer hallmark. While genetic alterations linked to cancer metabolism are the focus of intense research, the role of non-coding RNAs such as microRNAs, long non-coding RNAs and circular RNAs as initiators of cancer metabolic alterations is poorly studied. In the present manuscript we unveil the impact of circPVT1 aberrant expression on breast cancer metabolism. Indeed, we originally found that circPVT1 sponges the metabolic miRNA-33a-5p, thereby releasing aberrantly c-MYC transcriptional activity that leads to deregulated expression of glutaminase (Fig. 6l). Despite to a less extent than miR-33a-5p, we found, under the same experimental conditions, that circPVT1 sponges miR-145-5p and miR-203 (Fig. S1e-h). Being c-MYC a well-known target of all three microRNAs this might suggest that circPVT1 tightly controls c-MYC expression by sponging either specifically or concomitantly specific microRNAs. Both circPVT1 sponging specificity and concomitance on miR-33a-5p or other microRNAs could be dictated either by the tumoral genomic landscape or by the stage of a given tumorigenic process. There is growing evidence that metabolic alterations represent a rapid and adaptive response to oncogenic cues that might occur independently from genetic alterations. Thus, the formation of specific circRNA/microRNAs interacting codes, as herein described, might confer spatial and temporal plasticity to the metabolic adaptation of breast cancer cells.

Glutamine is a fundamental amino acid for many functions of cancer cells. It has been reported that glutamine consumption is a very active process in cancer cells when compared to that of other amino acids. TNBC subtype which lacks oestrogen receptor (ER) expression, progesterone receptor (PR) expression, Epidermal Growth factor receptor 2 (HER2) has been reported to be more glutamine addicted than other breast cancer subtypes. At which stage of breast tumorigenic process the role of these non-coding interacting codes could preferentially be exerted, is still underexplored. One of the unmet clinical needs in daily cancer treatment is represented by the relapse of the primary tumor either locally, skin metastasis or at distant sites. Growing evidence document that frequently distant metastasis and matched primary tumors including TNBC exhibit a rather similar mutational landscape. These findings imply that additional non-genetic alterations might contribute the to the establishment and chemoresistance of the metastatic phenotype. Among them glutamine addiction fuelled by aberrant non-coding RNAs network as that involving circPVT1/miR-33a-5p which unleashes c-MYC-GLS1 transcriptional axis might play an important role. While the pivotal role of c-MYC in cancer metabolic alteration has been established, the mechanistic events governing its aberrant activity have not been fully explored. Herein, our findings identify an upstream post-translational network leading to MYC accumulation and transcriptional activity.

There is growing evidence on the therapeutic potential of inhibiting glutamine metabolism to treat human cancers. This attempt has been so far quite challenging. Indeed, the use of specific GLS1 inhibitors has been very successful in preclinical cellular systems while it has been poorly efficacious in mouse models of pancreatic cancer [25]. Recent evidence documents that broad inhibition of glutamine metabolism rather than inhibition of glutaminolysis might hold higher therapeutic potential. 6-diazo-5-oxo-L-norleucine (DON) has anticancer effects but its therapeutic application has been hampered by gastrointestinal toxicity [25]. Recently, DON has been shown to impair the growth of both primary and metastatic PDA in PDOs and transplantable models. We found that the GLS1 inhibitor BPTES exerted a dose-dependent anti-tumoral activity on PDOs derived from three TNBC metastatic patients (Fig. 6f). Recently, a first-in human biomarker-driven Phase I trial (NCT03894540) testing selective inhibitors of GLS1 (IACS-6274) in patients with solid tumors has been concluded. The results of the trial document that IACS-6274 is well tolerated, with good PK, significant target modulation and preliminary anti-tumoral activity [26]. In aggregate these findings illustrate a scenario in which the choice of the therapeutic targeting of either glutaminolysis or glutamine metabolism might be related to type of the cancer, to its stage as primitive tumor or metastatic disease, and to previously administered cancer treatment. Furthermore, the molecular events occurring transcriptionally or post-transcriptionally and dictating the alteration of glutaminolysis or having a broader impact on glutamine metabolism could also play a major role. It is still uncertain at which stage along a given tumorigenic process aberrant GLS1 activity takes place. The elucidation of temporal and spatial GLS1 activity might contribute to design combinatorial treatment targeting broadly glutamine metabolism.

## List of abbreviations

miRNAs: micro RNAs
circRNA: circular RNAs
lncRNA: long non-coding RNAs
GLUD1: glutamate dehydrogenase 1
TNBC: Triple negative breast cancer
TCA: Tricarboxylic acid cycle
GLS1: Glutaminase 1
PDO: Patient derived organoid
BC: Breast cancer
TCGA: The Cancer Genome Atlas
ChIP: Chromatin immunoprecipitation
ER: estrogen receptor
PR: progesterone receptor
HER2: Epidermal Growth factor receptor 2
DON: 6-diazo-5-oxo-L-norleucine
PK: Pharmacokinetics

## Acknowledgements

Not applicable

## Authors’ contributions

AC.P., C.P., performed the experiments and interpreted the data; D.R., S.D., and C.B generated PDO; A.S. performed the bioinformatics analyses; V.D.P. performed nucleic acid extraction and digital PCR. L.C. and MC. V. performed H-NMR analysis and interpretated the related data; R.F.A. performed seahorse metabolic analysis; AC.P., C.P., S.S., G.B. wrote the manuscript. P.M., F.P. critically revised the manuscript. S.S and G.B conceived and supervised the research activities.

## Ethics approval and consent to participate

Generation of patient-derived organoids from breast cancer was approved by institutional review board of Regina Elena National Cancer

## Consent for publication

Not applicable.

## Declarations

Institute and appropriate regulatory authorities (approval no. IFO 1270/19). All patients signed an informed consent.

## Availability of data and materials

All data supporting the conclusions of this article are included within the article and its additional files. Any other information is available from the corresponding authors upon reasonable request.

---

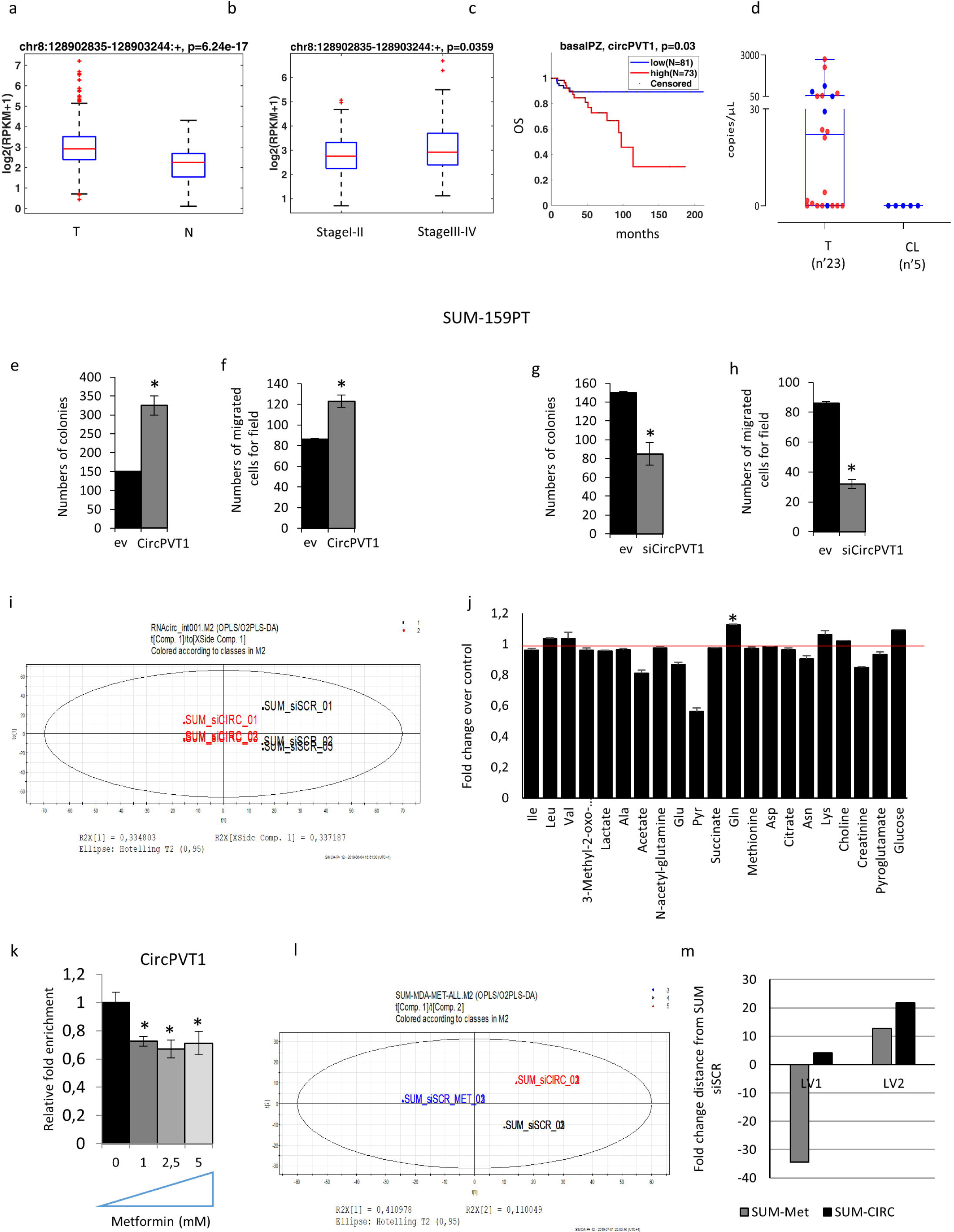

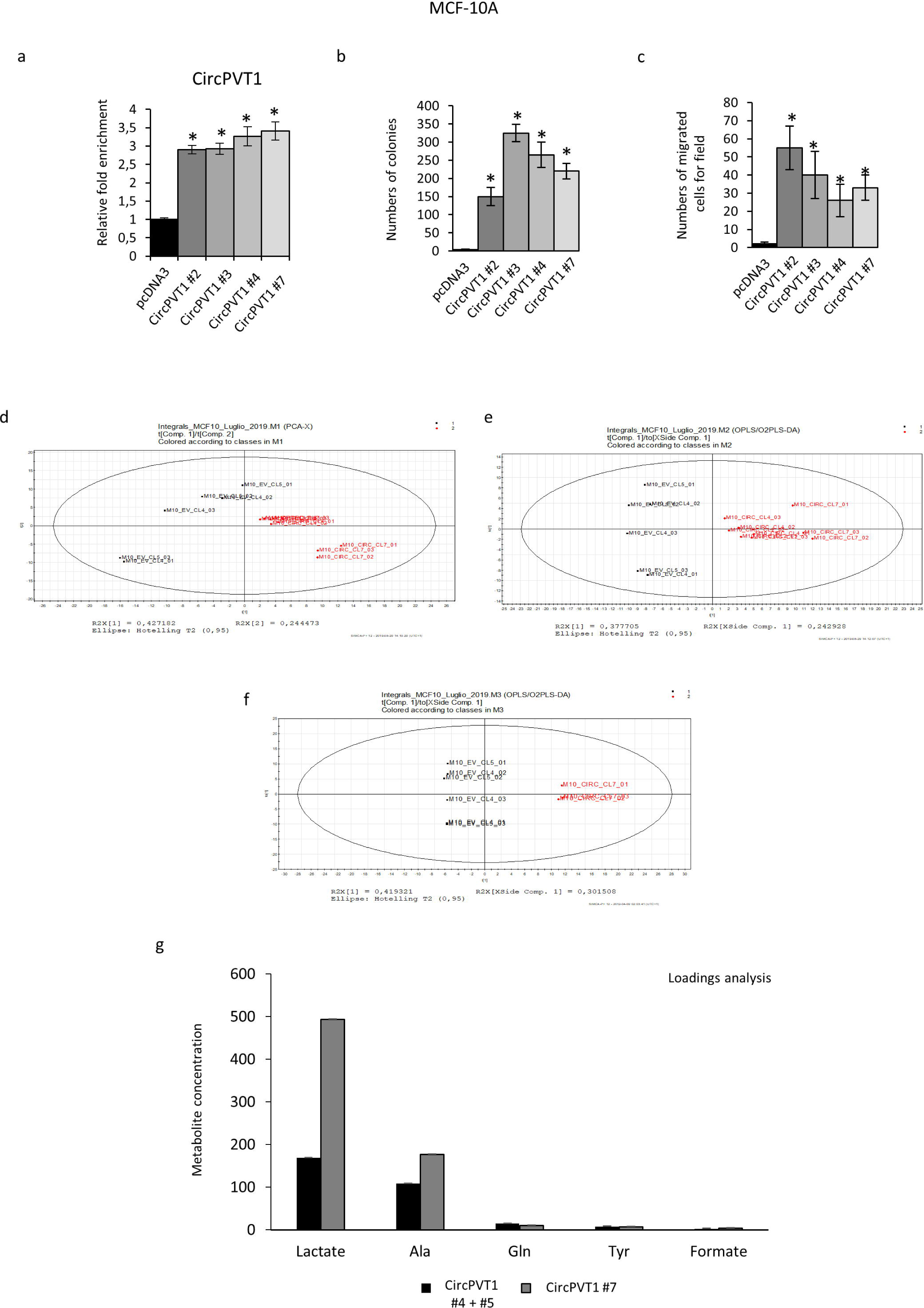

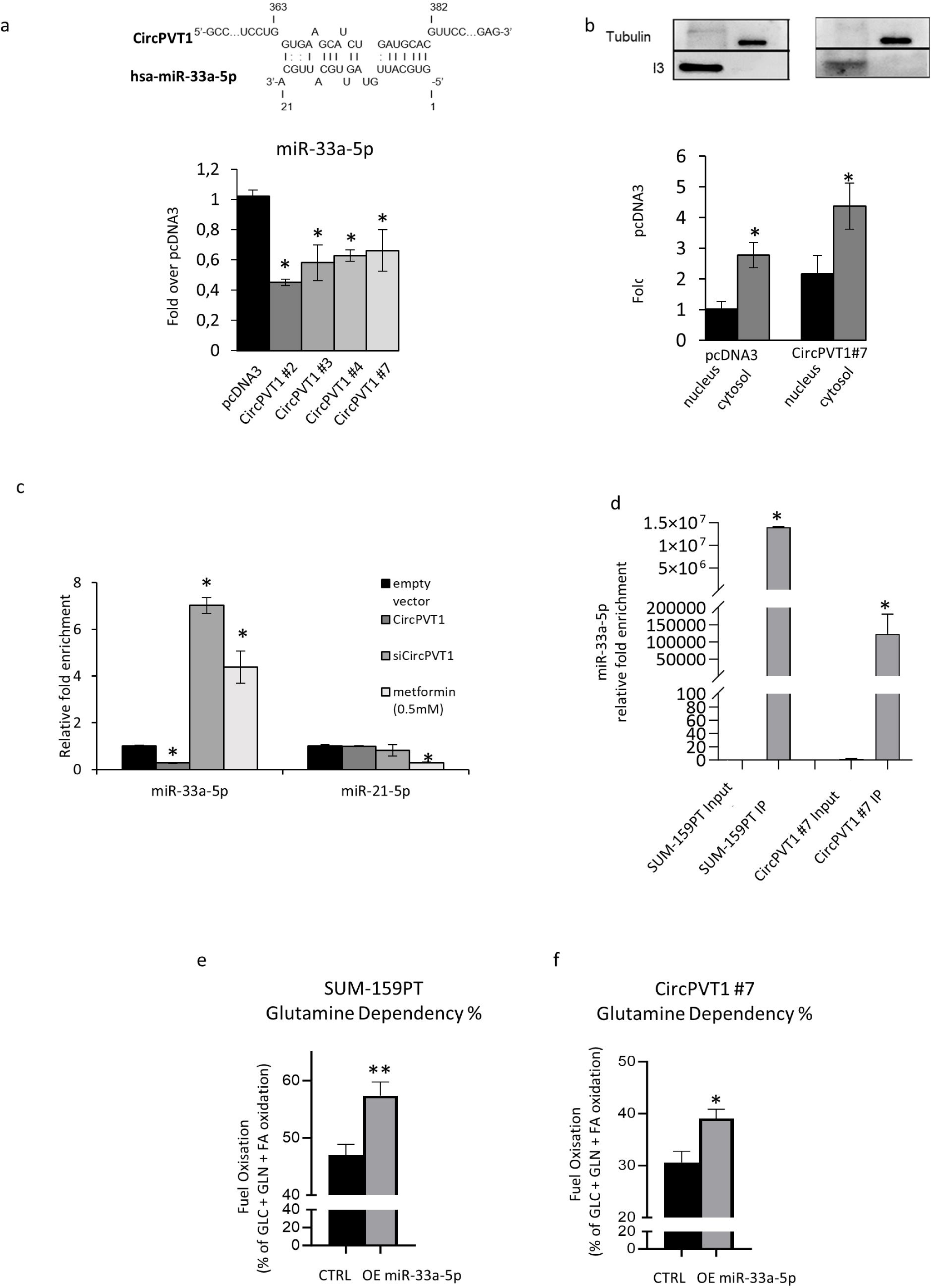

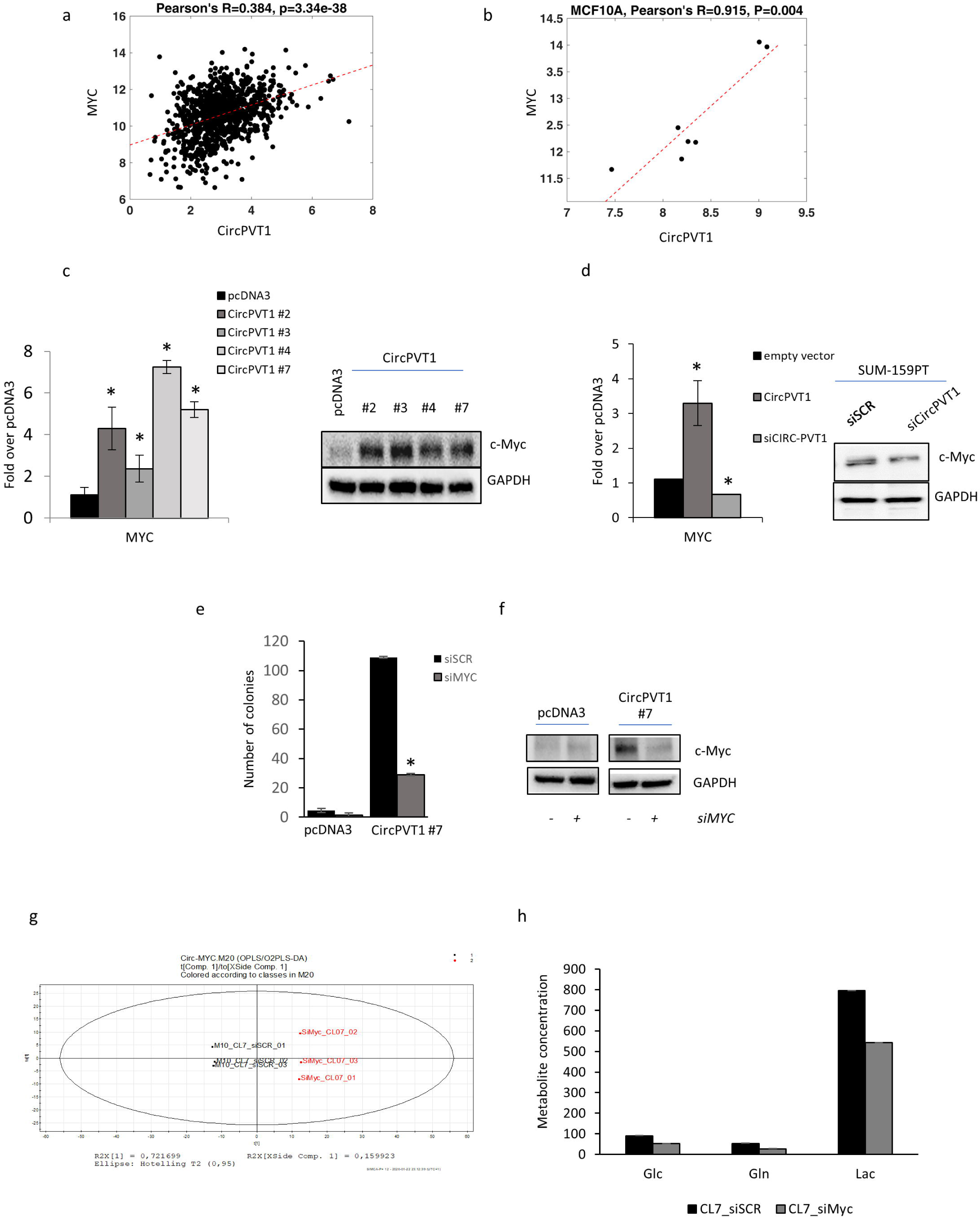

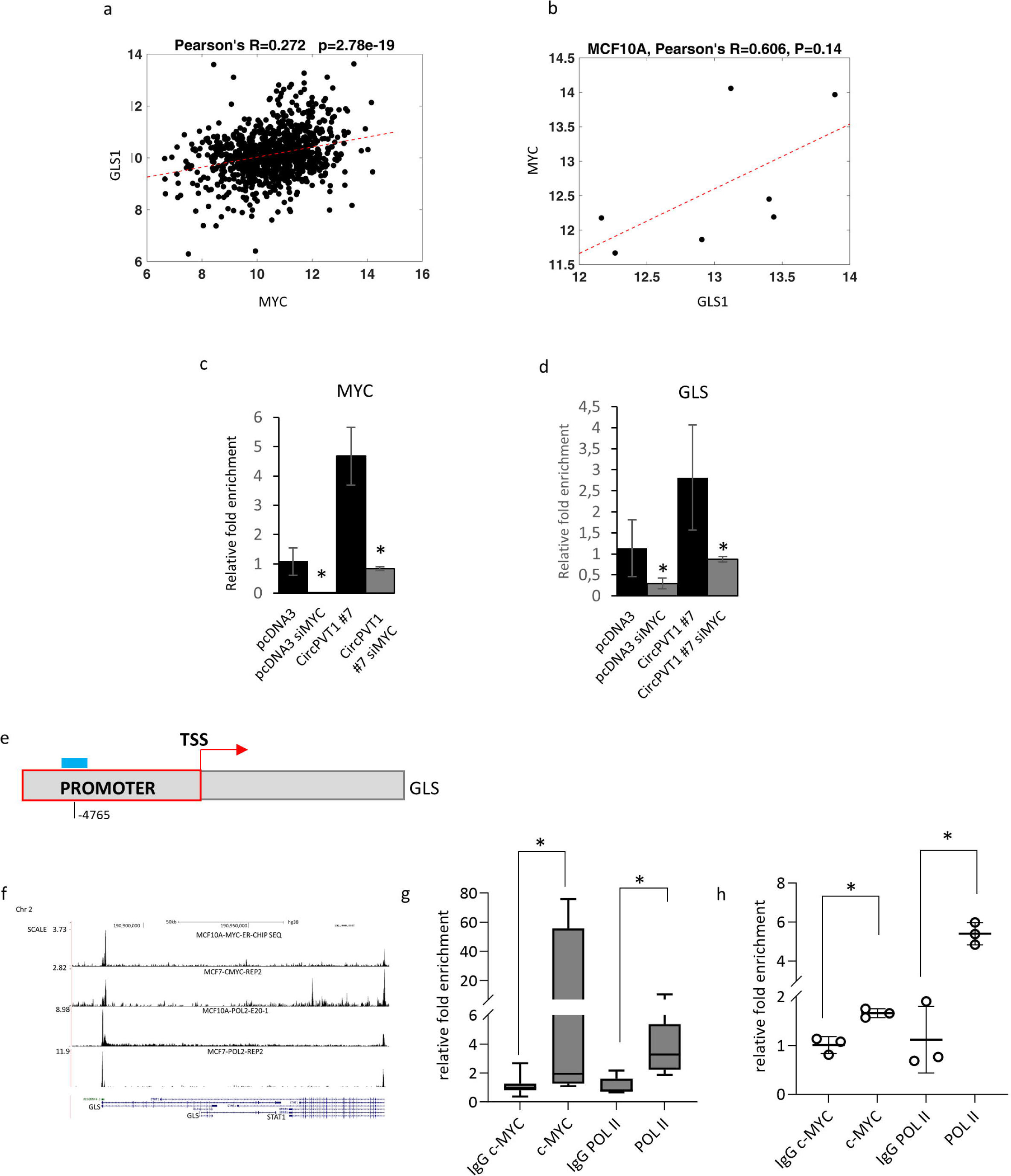

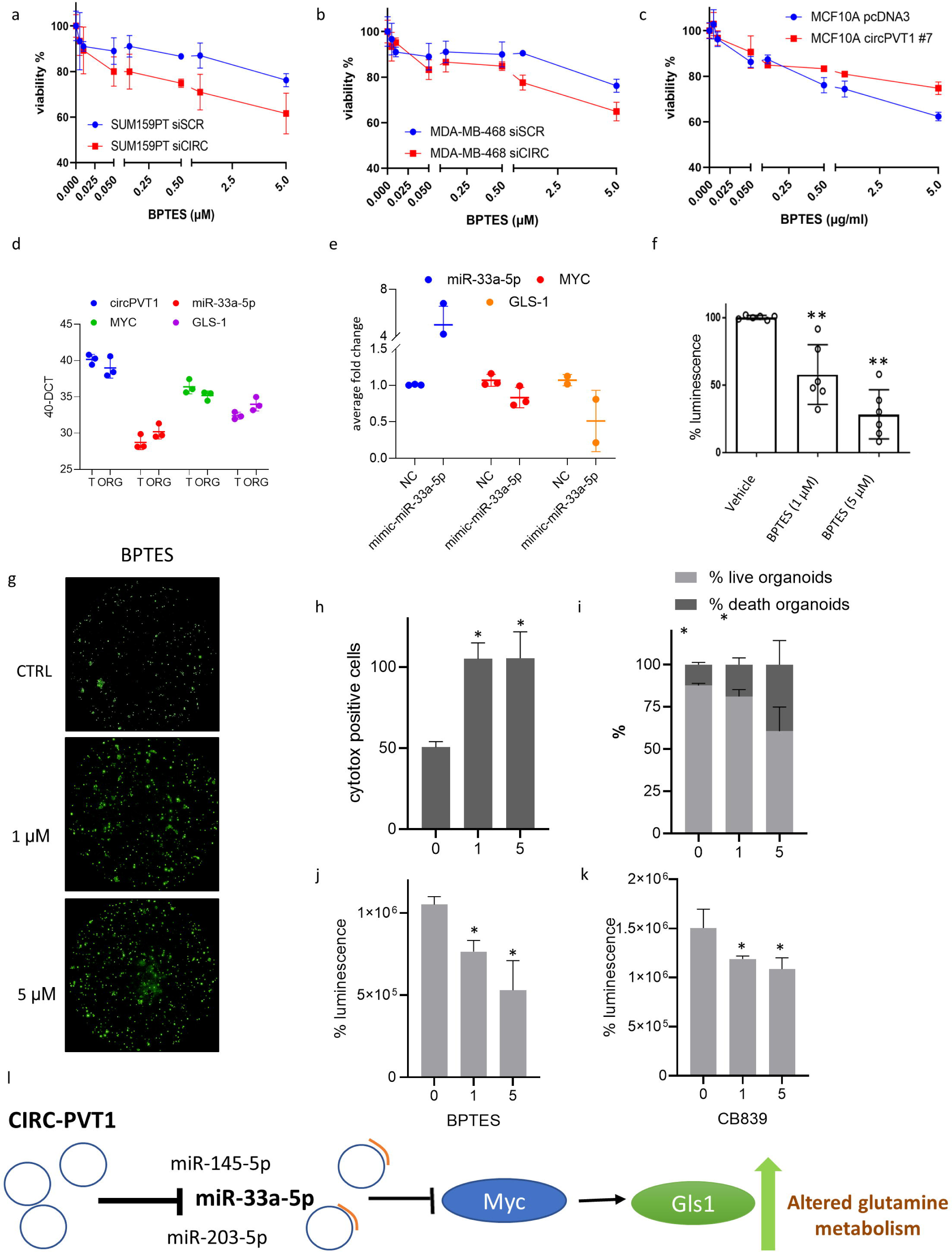

