## Supplementary figures and images for "CircPVT1 sponges miR-33a-5p unleashing the c-MYC/GLS1 metabolic axis in breast cancer"

### Supplemental Figure 1

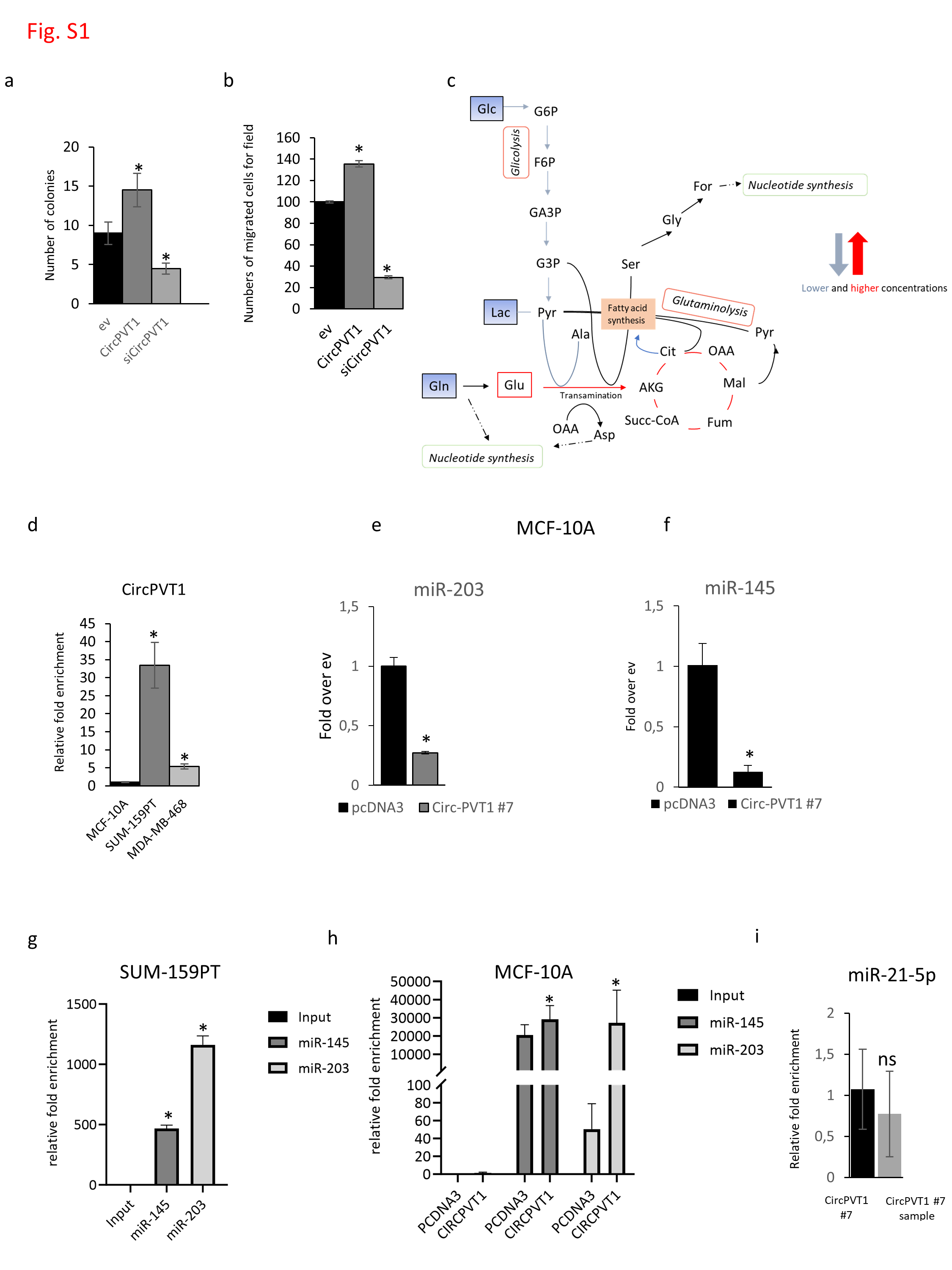

### Supplemental Figure 2

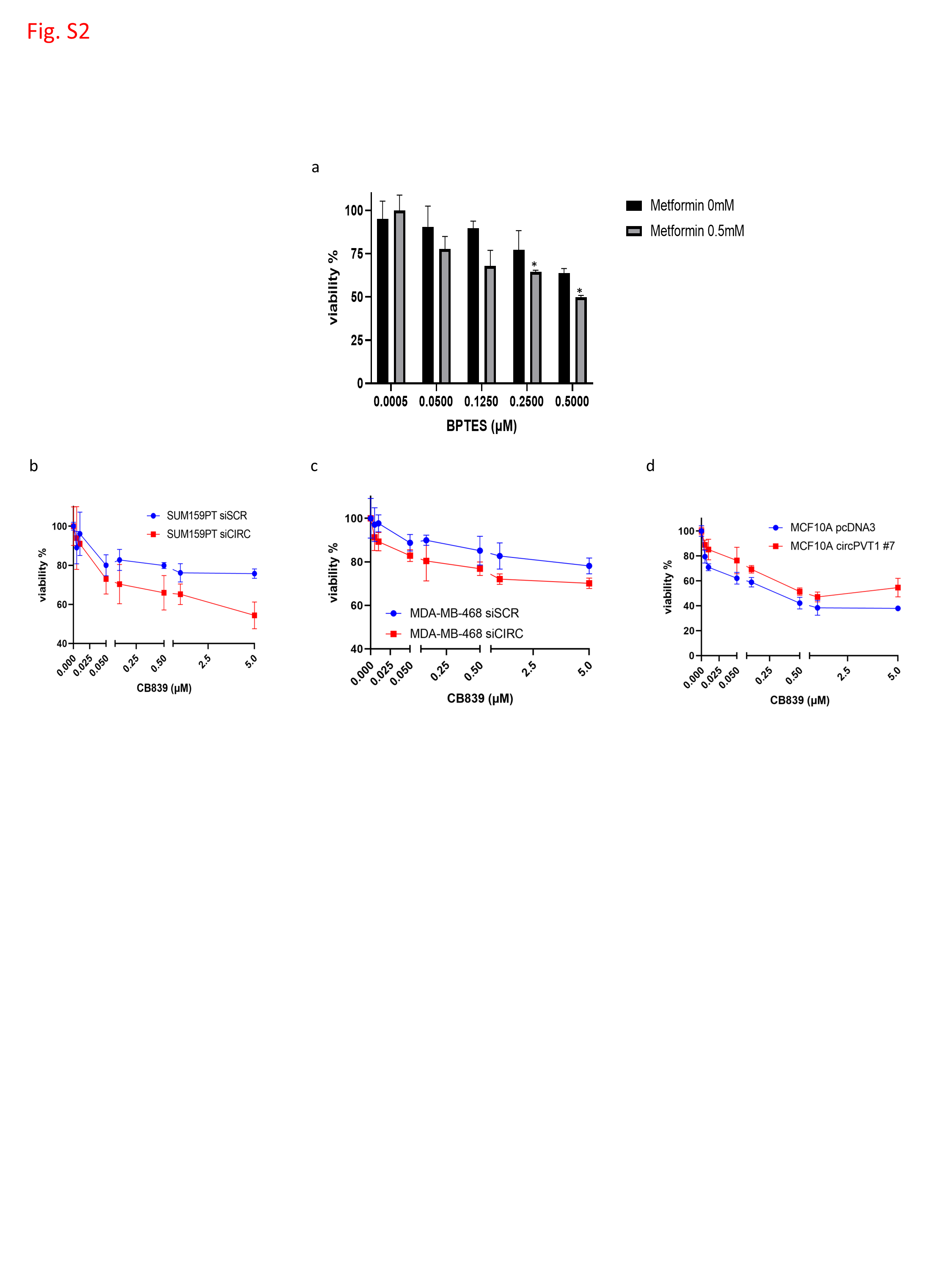
