## Supplemental Figure Legends for "CircPVT1 sponges miR-33a-5p unleashing the c-MYC/GLS1 metabolic axis in breast cancer"

**Fig.1** **a-b** Log2 expression levels of chromosome interval containing circPVT1 in tumor and non-tumoral samples (a) and in stage I-II and III-IV tumor samples (b). **c** Kaplan Meier curves indicating the overall survival of patients basing on the expression level of circPVT1. **d** Boxplots show the copies/μl of circPVT1 in 23 TNBC tumors and in 5 controlateral breast tissues. **e-g** Histograms show the number of colonies of SUM-159PT cells either expressing high (e) or low levels (g) of circPVT1. **f-h** Histograms show the number of migrated SUM-159PT cells treated as in f-g. **i** PCA models built on the 1H-NMR dataset of media samples cell extracts from SUM-159PT cell cultures either expressing endogenous or low levels of circPVT1. **j** Histograms show the fold changes of the most discriminant metabolites between the two groups from the PCA models (i). **k** Relative fold enrichment of circPVT1 levels measured in SUM-159PT cells treated with increased concentration of Metformin. **l** PCA models built on the 1H-NMR dataset of media samples cell extracts from SUM-159PT cell cultures either expressing endogenous levels or silenced for circPVT1 and treated or not with 0.5 mM of metformin. **m** t-test applied on LV1 and LV2 component on SUM_siSCR treated with metformin and SUM_sicircPVT1.

**Fig.2** **a** CircPVT1 RNA relative enrichment levels measured in MCF-10A stably expressing pcDNA3 or circPVT1. Numbers (#) indicated the different clones of MCF-10A stably expressing high circPVT1 levels. **b-c**. Histograms show number of colonies (b) or of migrated cells (c) count in MCF-10A treated as in a. **d-e** PCA (d) and OPLS-DA (e) models built on the 1H-NMR dataset of media samples cell extracts from MCF-10A cell cultures (clones #2, 4 and 7) either expressing endogenous or high levels of circPVT1. **f** OPLS-DA model built on the 1H-NMR dataset of media samples cell extracts from MCF-10A cell clone #7 either expressing endogenous or high levels of circPVT1. **g** Histograms show the fold changes of the most discriminant metabolites between the different groups from the PCA model.

**Fig.3** **a** Upper part, predictive site of binding interaction between miR-33a-5p and circPVT1. Lower part, histograms show the expression levels of miR-33a-5p in MCF-10A cells treated as in Figure 2a. **b** Upper part, representative protein gel blot of nucleus/cytosol cell lysates obtained from MCF-10A cells stably expressing pcDNA3 or circPVT1 (clone #7) stained with the indicated antibodies. Lower part, histograms show the levels of circPVT1 between nucleus/cytosol obtained from MCF-10A treated as in the upper part. **c** Histograms show the expression levels of miR-33a-5p and miR-21-5p measured in SUM-159PT expressing endogenous or high or low levels of circPVT1 or treated with 0.5 mM of metformin. **d** Histograms show the miR-33a-5p relative fold enrichment in SUM-159PT and MCF-10A circPVT1#7 cells measured in total RNA immunoprecipitated with circPVT1-capture probes. **e-f** Histograms show the fuel oxidation rate of SUM-159PT (e) and MCF-10A circPVT1#7 cells (f) expressing endogenous or ectopic levels of miR-33a-5p.

**Fig.4** **a-b** Pearson positive correlation between MYC and circPVT1 expression levels from TCGA breast data set (a) or from seven different clones of MCF-10A ectopically expressing high levels of circPVT1 (b). **c** Histograms show the MYC expression levels measured in MCF-10A treated as in Figure 2a and representative protein gel blot of whole cell lysates extracted from MCF-10A treated as in Figure 2a. **d** Histograms show the MYC expression levels measured in SUM-159PT expressing endogenous or high or low levels of circPVT1 and representative protein gel blot of whole cell lysates extracted from SUM-159PT cells silenced or not for circPVT1. **e** Histograms show the number of colonies of MCF-10A clone #7 after c-Myc silencing. **f** Representative protein gel blot of whole cell lysates extracted from MCF-10A cells silenced or not for c-Myc. **g** PCA models built on the 1H-NMR dataset of media samples cell extracts from MCF-10A cell cultures either expressing endogenous or stably expressing high levels of circPVT1 followed silencing of MYC or SCR. **h** Histograms show the fold changes of the most discriminant metabolites between the four groups from the PCA models (g).

**Fig.5** **a-b** Pearson positive correlation between MYC and GLS expression levels from TCGA breast data set (a) or from seven different clones of MCF-10A ectopically expressing high levels of circPVT1 (b). **c-d** Histograms show the relative MYC (c) and GLS (d) expression levels measured in MCF-10A pcDNA3 and circPVT1 #7 silenced for SCR or MYC. **e** Predictive binding region of c-Myc on GLS promoter region. **f** c-Myc protein enrichment on GLS promoter region in MCF-10A cells from ChIP-seq data deposited on the CistromeDataBase (http://cistrome.org/). **g-h** Relative enrichment of the occupancy of c-Myc p-Ser62 on the regulatory regions of GLS (-4850: -4811 upstream GLS transcriptional starting site) assessed by Chromatin Immunoprecipitation in MCF-10A circPVT1 #7 (g) or SUM-159PT (h).

**Fig.6 a** Viability curves obtained by measuring ATP levels in SUM-159PT cells silenced or not for circPVT1 and treated for 72 hrs with increasing doses of BPTES (0 – 5 µM). **b** Viability curves obtained by measuring ATP levels in MDA-MB-468 cells silenced or not for circPVT1 and treated for 72 hrs with increasing doses of BPTES (0 – 5 µM). **c** Viability curves obtained by measuring ATP levels in MCF-10A circPVT1#7 and treated for 72 hrs with increasing doses of BPTES (0 – 5 µM). **d** Relative fold enrichment of circPVT1, miR-33a-5p, MYC and GLS among tumoral and matched patients derived organoids (ORG) (n=3). **e** Relative fold enrichment of miR-33a-5p, MYC and GLS1 following miR-33a-5p overexpression. **f** Histograms show PDOs (n=3) viability after 72 hrs of BPTES treatment (1 and 5 µM). **g-h** Cytotox positive cells of #240 after 72 hrs of BPTES treatment at different doses (1 and 5 µM). **i** Percentage of lived organoids over death organoids after #240 treatment as in G-H. **j** Percentage of luminescence of #240 organoids treated as in g-h. **k** Percentage of luminescence of #240 organoids after 72 hrs of CB839 treatment at different doses (1 and 5 µM). **l** CircPVT1 alters glutamine metabolism by sponging metabolic miR-33a-5p, sustaining c-Myc translation that in turn promotes GLS transcription.
